# MOSCATO: A Supervised Approach for Analyzing Multi-Omic Single-Cell Data

**DOI:** 10.1101/2021.09.02.458781

**Authors:** Lorin M Towle-Miller, Jeffrey C Miecznikowski

## Abstract

Advancements in genomic sequencing continually improve personalized medicine in complex diseases. Recent breakthroughs generate multiple types of signatures (or multi-omics) from each cell, producing different data ‘omic’ types per single-cell experiment. We introduce MOSCATO, a technique for selecting features across multi-omic single-cell datasets that relate to clinical outcomes. For example, we leverage penalization concepts often used in multi-omic network analytics to accommodate the high-dimensionality where multiple-testing is likely underpowered. We organize the data into multi-dimensional tensors where the dimensions correspond to the different ‘omic’ types. Using the outcome and the single-cell tensors, we perform regularized tensor regression to return a variable set for each ‘omic’ type that forms the clinically-associated network. Robustness is assessed over simulations based on available single-cell simulation methods. Real data comparing healthy subjects versus subjects with leukemia is also considered in order to identify genes associated with the disease. The flexibility of our approach enables future extensions on distributional assumptions and covariate adjustments. This algorithm may identify clinically-relevant genetic patterns on a cellular-level that span multiple layers of sequencing data and ultimately inform highly precise therapeutic targets in complex diseases. Code to perform MOSCATO and replicate the real data application is publicly available on GitHub at https://github.com/lorinmil/MOSCATO and https://github.com/lorinmil/MOSCATOLeukemiaExample.

## 1 Introduction

Classic bulk genetic sequencing involves averaging signature levels across all cells. Different sequencers may sequence different types of molecules such as ribonucleic acid (RNA), proteins, DNA methyl groups, etc. Disease progression, therapy success, and other clinical outcomes often vary among individuals suffering from complex diseases [3, 7, 17, 32], and the heterogeneity in their outcomes may be better understood through the intricacies of a patient’s molecular signatures [5, 25, 24, 1]. This has led to an explosive demand for multi-omics which involves integrating multiple types of molecular information in order to have a more Systems Biology approach. For example, in breast cancer patients with resistance to lapatinib therapy, Komurov et al. were able to suggest additional therapy targets by identifying combinations of RNA and proteins responsible for glucose deprivation that was associated with the resilience [18].

Methods for identifying graphs and gene regulatory networks within a single molecular type has been well studied [20, 14], however, different methods should be considered when integrating multiple types of molecular information in order to accommodate the between and within molecular relationships [6]. Each molecular type often contains thousands of features, and integrating them creates a higher dimensional problem with more sophisticated relationships both within and between molecular types. For example, the Decomposition of Network Summary Matrix via Instability (DNSMI) method decomposes a matrix of network strengths by fitting a series of models for the expected relationships across molecular types and with the disease outcome [35]. Supervised sparse Canonical Correlation Analysis (SCCA) attempts to optimize the correlation matrix between molecular types through lasso constrained linear combinations of the features and also eliminates features weakly correlated with the outcome [33].

In bulk sequencing experiments, rare cells or smaller cell-types will be diluted due to the averaging across all cells within the sample. This motivated single-cell sequencing techniques where molecular information could then be sequenced on a cell-by-cell basis. While initial protocols were limited to RNA [23, 26], newer technology may now sequence multiple types of molecular information within each cell, denoted as *multi-modal* (or *multi-omic*) single-cell sequencing. For example, CITE-seq simultaneously sequences both cell surface proteins and RNA on each cell of a sample [29]. Although still a growing technology, applications have already been considered using this novel sequencing approach. For example, Kendal et al. utilized CITE-seq technology to compare tendons in healthy individuals to those with tendinopathy [15]. Figure 1 displays an example of single-cell data from each patient.

**Figure 1:**
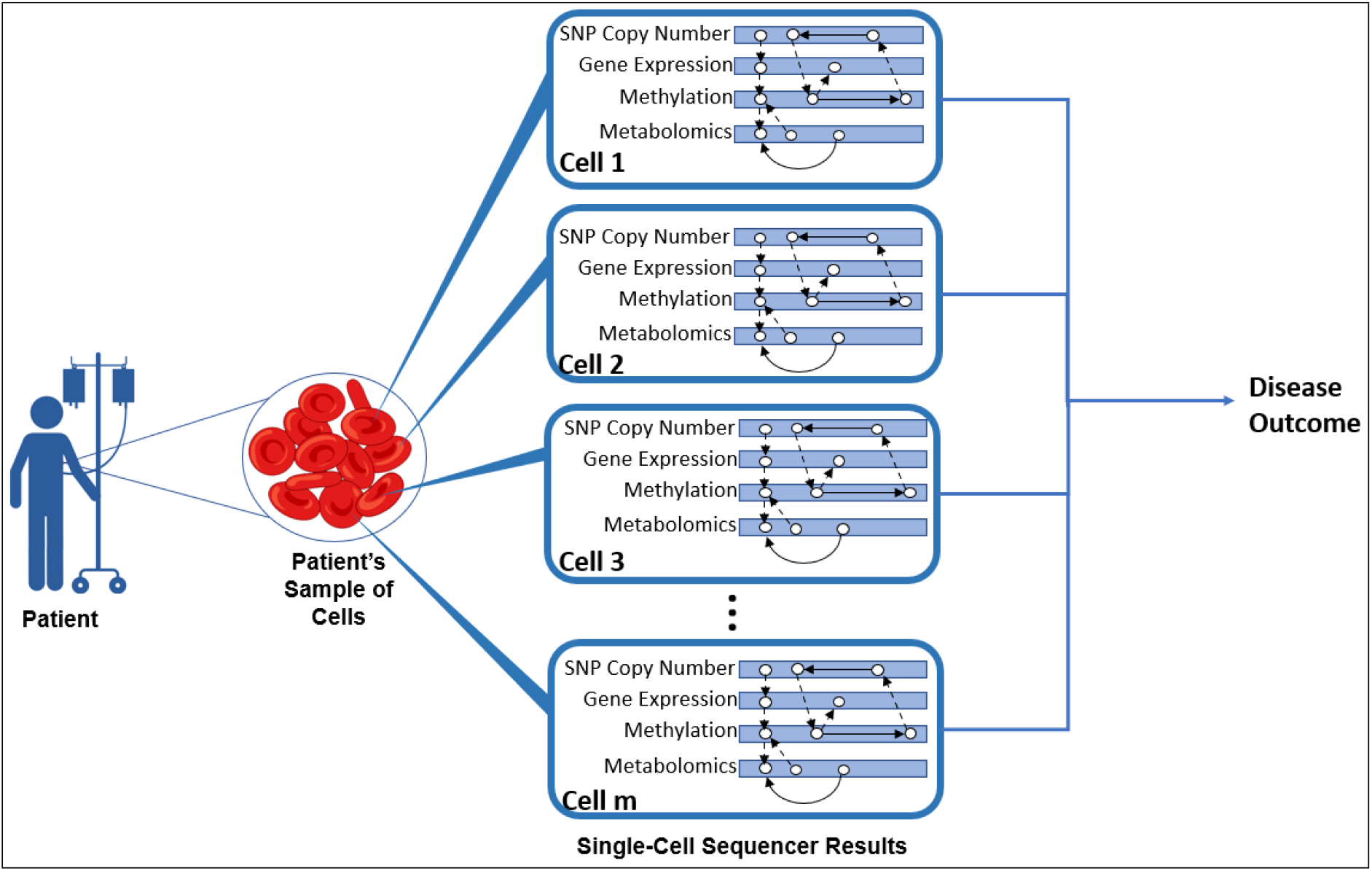
Pictoral demonstration of single-cell network detection experiments and studies.

This manuscript proposes a novel method, Multi-Omic Single-Cell Analysis using TensOr regression (MOSCATO), for identifying the superstructure of a semi-directed graph, or *network*, within multi-omic single-cell data that relates to a disease or phenotypic outcome. Section 2 describes preliminary tensor concepts, Section 3 introduces MOSCATO, Section 4 performs simulations of multi-omic single-cell data and applies MOSCATO under various scenarios, Section 5 applies MOSCATO to real single-cell data, and Section 6 discusses future work and limitations.

## 2 Preliminaries

MOSCATO utilizes regularized tensor regression, and this section describes existing and relevant tensor concepts. Section 2.1 defines tensors and basic tensor operations, and Section 2.2 uses the operations and definitions from Section 2.1 to describe tensor regression and regularization techniques.

### 2.1 Tensor Definitions

High dimensional data may be organized into a tensor, and a matrix may be thought of as a 2-dimensional tensor. Utilizing familiar tensor notation as provided by Kolda and Bader [16], we let 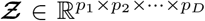 denote a D-dimensional tensor where dimension *d* contains *p*_*d*_ variables for *d* = 1, …, *D*. For example, *D* = 1 denotes a vector and *D* = 2 denotes a matrix. Many mathematical operations for tensors build on mathematical operations used in matrices. For example, Definition 2.1 describes *outer products* between D vectors to create a D-dimensional tensor, where ∘ denotes the *Khatri-Rao product*.

#### Definition 2.1

*Let* ***b***_**1**_, ***b***_**2**_, …, ***b***_***D***_, *denote vectors where* 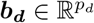. *Then the* ***outer product*** *of those vectors*, ***b***_**1**_ ∘ ***b***_**2**_ ∘ … ∘ ***b***_***D***_, *creates a D-dimensional tensor of size p*_1_ × *p*_2_ × … × *p*_*D*_, *and each* (*i*_1_, …, *i*_*D*_)^*th*^ *element equals* 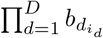.

It may also be convenient to reorganize a tensor into a lower dimensional space by *vectorizing* or *mode-d matricizing* the tensor. Definitions 2.2 and 2.3 describe these reorganization techniques.

#### Definition 2.2

*Let* 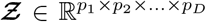 *denote a D-dimensional tensor. Then* 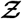 *may be reorganized into a column vector through the* ***vec*** *operator* 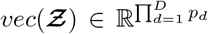, *where the* 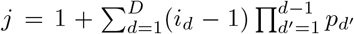 *element of vec*(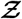) *corresponds to the* (*i*_1_, …, *i*_*D*_)^*th*^ *value in* 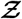.

#### Definition 2.3

*Let* 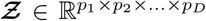 *denote a D-dimensional tensor. Then* 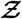 *may be reorganized into a matrix through the* ***mode-d matricization*** *operator* 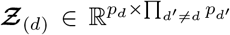, *where the* (*i*_*d*_, *j*)^*th*^ *element within* 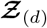 *equals the* (*i*_1_, …, *i*_*D*_)^*th*^ *value within* 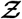 *and j* = 1 + Σ_*d*′≠*d*_(*i*_*d*′−1_)Π_*d*′′<*d*′,*d*′′≠*d*_ *p*_*d*′′_.

Similarly as done in matrix operations, it may be of interest to multiply two tensors with comparable dimensions via *inner products*, as described in Definition 2.4.

#### Definition 2.4

*Suppose two tensors* 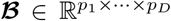 *and* 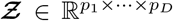. *The* ***inner product*** *may be obtained by*

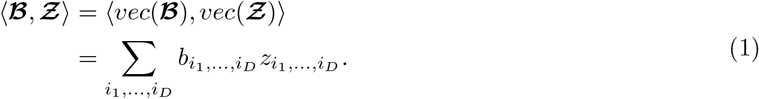

Furthermore, it may be of interest to multiply a matrix along the *d*^*th*^ dimension of a tensor through *d-mode products* as described in Definition 2.5.

#### Definition 2.5

*Suppose a tensor* 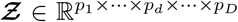 *and a matrix* 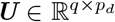. *The* ***d-mode product*** *between* 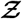 *and* ***U*** *may be expressed as* 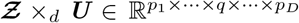 *where the* (*i*_1_, …, *i*_*d*−1_, *j, i*_*d*+1_, …, *i*_*D*_)^*th*^ *value equals* 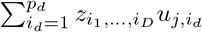.

The rank of a matrix denotes the maximum number of linearly independent rows/columns in the matrix. Building on those concepts, the rank of a tensor may be thought of as the maximum number of vectors that can be multiplied and added to replicate the tensor, as shown in Definition 2.6.

#### Definition 2.6

*Assume a D-dimensional tensor* 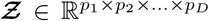. 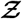 *is rank-R if there exists vectors* 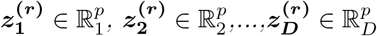 *for r* = 1, 2, …, *R such that*

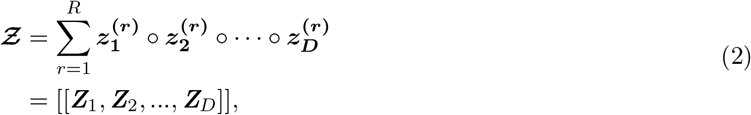

*where* 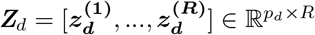.

The true rank of a tensor may often be difficult to determine due to the high dimensionality, motivating decomposition techniques that estimate vectors for a given rank that approximate the tensor, as shown in Definition 2.7.

#### Definition 2.7

*Assume a D-dimensional tensor* 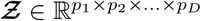. *A* ***rank-****R* ***CP decomposition*** *aims to use R vector sets (one vector per dimension) to approximate* 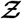 *by*

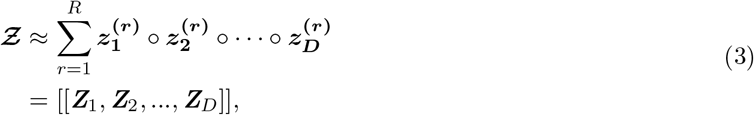

*where* 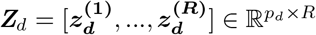.

Kolda and Bader present additional details on decomposition and other tensor operations [16].

### 2.2 Tensor Regression

Building on notation covered in Section 2.1, this section will briefly describe tensor regression that was originally presented by Zhou et al. [36]. Tensor regression builds on Generalized Linear Model (GLM) concepts, where we have an outcome *y* for each subject that follows some exponential family with link function *g*(·) and mean *µ*. Classic GLM uses univariate independent variables to predict the outcome, but tensor regression extends those concepts by additionally allowing a predictor tensor. This is accomplished by multiplying the dimensions of the tensor through coefficient vectors that convert the dimensions to a common univariate value that may then predict the outcome.

Figure 2 shows a simple example with rank-1 tensor regression and *D* = 2 dimensions for the predictor tensor. Each dimension in the predictor tensor corresponds to a different feature set, and each subject will contain its own *D* = 2 predictor tensor to be used to predict their univariate outcome *y*.

**Figure 2:**
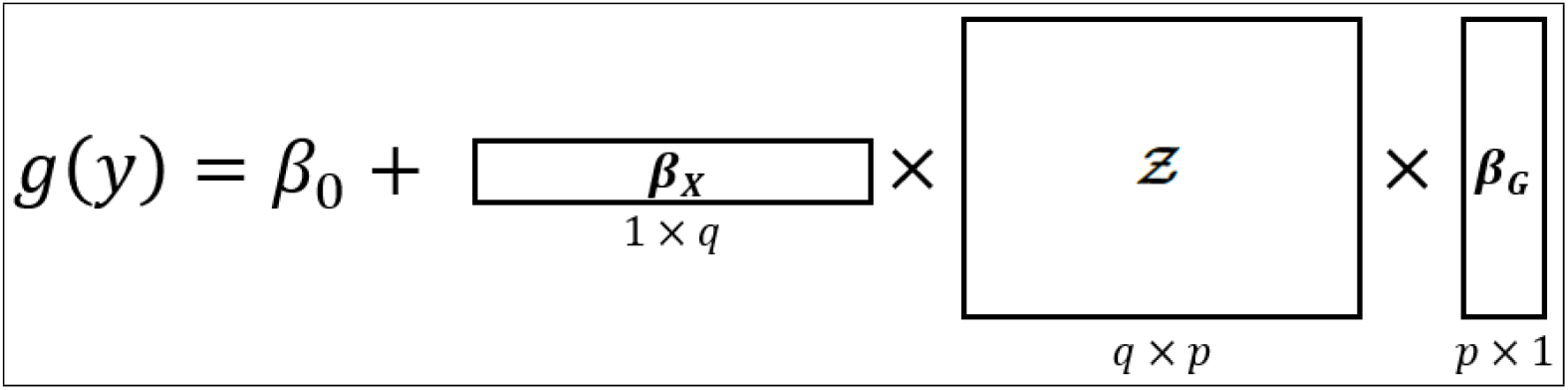
Simple Example of Rank-1 Tensor Regression with *D* = 2. Suppose a univariate outcome *y* with canonical link *g*(·) and predictor tensor 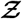 with *D* = 2 dimensions. Each dimension in the predictor tensor corresponds to a feature set, and each feature set contains its own coefficient vectors. The coefficient vectors in this example, ***β***_***X***_ and ***β***_***G***_, may be estimated by collecting outcomes and predictor tensors across multiple subjects and applying the Block Relaxation Alogorithm.

A motivating example for when these types of models may be useful is when each subject contains their own magnetic resonance imaging (MRI) image of their brains, and suppose each subject has an outcome specifying their disease status (e.g., brain tumor versus no brian tumor). These images may be expressed as a dataset (i.e., a 2-dimensional tensor) by organizing the images into compably sized grids where each grid point denotes a pixel from the image, and the value within each pixel quantifies the amount of pigment from the image. Referring to the model shown in Figure 2, the MRI image would correspond to the predictor tensor, 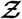, and coefficients would be estimated such that by inputting a subject’s MRI image could then predict whether they had a brain tumor. The coefficient vectors in this example could help describe which regions of the brain predict whether someone has a brain tumor (i.e., which rows/columns contain high/low coefficient values).

Tensor regression may also involve higher rank problems with the more formal representation

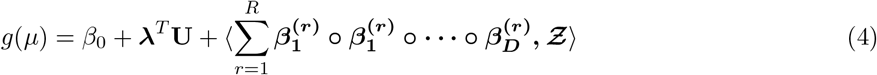

where **U** contains the univariate independent variables. A rank-*R* tensor regression estimates *R* coefficient vectors for each dimension in the predictor tensor, but for simplicity, in this manuscript we will assume rank-1. The *Block Relaxation Algorithm* is used to estimate the coefficient vectors with additional details described by Zhou et al. [36]. Zhou et al. [36] also claim that regularization in tensor regression may be accomplished by simply imposing constraints when fitting the models on each dimension.

If one naively vectorized the predictor tensor and fit a classic GLM model, it would require estimating 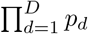 coefficients for the tensor. This approach would not only ignore the inherent structure of the data by treating each element in the tensor as independent with no distinction between the dimensions, it would also attempt to estimate many more coefficients compared to 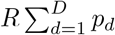 coefficients in tensor regression. Consequently, this naive approach may be unrealistic in high dimensional problems given a typically much smaller sample size. This reduction in parameters highlights the benefits of tensor regression. However, it is subject to limitations such as uniqueness and identifiability. For example, suppose a rank-1 model with 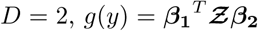. Then for any scalar *τ*, we could derive an equally optimal model 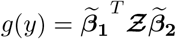 where 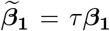 and 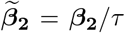. Additionally, the Block Relaxation Algorithm may converge to a local maxima as opposed to the global maxima when attempting to maximize the log likelihood. Measures to check for these concerns exist and are discussed in further detail by Zhou et al. [36].

## 3 Methods

### 3.1 The Model

In classic bulk sequencing, the data contains one record per subject. Supposing *n* subjects with two data types, bulk sequencing studies would contain two data sets (i.e., a dataset for each data type), ***𝒢*** ∈ ℝ^*n*×*p*^ and ***𝒳*** ∈ ℝ^*n*×*q*^. In single-cell sequencing, there are multiple records per subject where each row corresponds to a cell within the subject. Consequently, for a given subject *i* with two data types, their single-cell data would contain two datasets 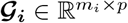 and 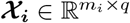, where *m*_*i*_ denotes the number of cells for subject *i*. Since the number of cells typically differs across subjects (i.e., *m*_*i*_ ≠ *m*_*i*′_ where *i* ≠ *i*′), we organize each subject’s data into separate datasets (i.e., ***𝒢***_***i***_ and ***𝒳***_***i***_) as opposed to organizing the input data directly into a 3-dimensional tensor with a dimension for cells. It should also be noted that each subject’s data often consists of thousands of cells, and concatenating the single-cell data in long format may be computationally inefficient. Furthermore, we assume each subject *i* contains a univariate outcome *y*_*i*_ for *i* = 1, …, *n*, and we may express the outcomes in a vector as **y** = [*y*_1_, *y*_2_, …, *y*_*n*_]^*T*^. For simplicity, we may denote the two data types as ***𝒢*** and ***𝒳*** without the *i* subscript, although as described previously, each subject’s single-cell data will contain separate matrices for the data types as opposed to expressing data in long format as found in bulk sequencing.

MOSCATO aims to identify a subset of features within ***𝒢*** and ***𝒳*** that relate to each other and the outcome. In graphical modelling terms, MOSCATO identifies the superstructure of a semi-directed graph with undirected nodes involving features within ***𝒢*** and ***𝒳*** with some path directed to the outcome **y**. MOSCATO accomplishes this by imposing elastic net constraints on a tensor regression model [37].

Similarly as in the MRI image example from Section 2.2, multi-omic single-cell data contains multi-dimensional data per subject (i.e., features within ***𝒢*** and features within ***𝒳***) with a univariate outcome. This motivates the use of tensor regression for multi-omic single-cell data. Additionally, tensor regression not only efficiently accommodates multi-dimensional input data with a univariate outcome, it also handles regularization techniques and allows for additional covariate adjustments (e.g., age, sex, race, etc.). However, tensor regression requires equivalent dimensions for each subject’s input tensor. Thus, to standardize the dimensions across each subject, the first step of MOSCATO involves estimating a correlation matrix between their data type matrices,

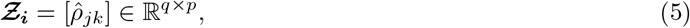

where 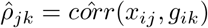 letting *x*_*ij*_ denote the *j*^*th*^ feature in ***𝒳***_***i***_ and *g*_*ik*_ denote the *k*^*th*^ feature in ***𝒢***_***i***_ for the *i*^*th*^ subject. Although many summary matrices could be considered such as the inverse of the covariance matrix or mutual information, Pearson correlation provides a simple interpretation while also standardizing the values within 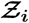 between -1 and 1.

Now using each of the 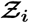 tensors for *i* = 1, …, *n* to estimate the coefficients, a tensor regression model similar to (4) and depicted in Figure 2 will be fit with elastic net constraints. The elastic net constraint works to balance by a weighted average between an *L*^1^-norm and *L*^2^-norm, where the *L*^1^-norm truncates small coefficients to zero and the *L*^2^-norm better handles highly correlated features. In summary, the elastic net constraint typically denoted as 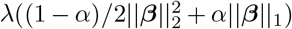 in the classical GLM setting will now involve

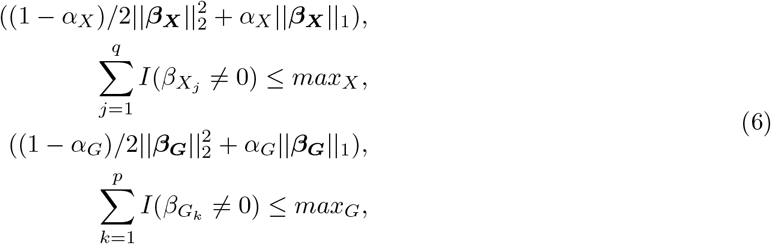

where ***β***_***X***_ ∈ ℝ^*q*^ denotes the coefficient vector for ***𝒳***, *β*_***G***_ ∈ ℝ^*p*^ denotes the coefficient vector for 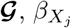 denotes the coefficient for the *j*^*th*^ feature in ***𝒳***, and 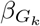 denotes the coefficient for the *k*^*th*^ feature in ***𝒢***. The hyperparameters *α*_*X*_ ∈ [0, 1] and *α*_*G*_ ∈ [0, 1] denote the weights to put on the *L*^1^-norm constraints for ***𝒳*** and ***𝒢***, respectively. The hyperparameter *λ* in the classical GLM setting denotes the overall weight to put on the constraint and it may be any positive number from 0 to infinity. Since tuning *λ* to the proper range may be difficult due to the nontrivial parameter space, we use *max*_*X*_ and *max*_*G*_ instead to denote the number of non-zero values within ***β***_***X***_ and ***β***_***G***_, respectively. This reparameterization of the constraints drastically simplifies the hyperparameter space and subsequent tuning described in Section 3.2.

For some fixed *α*_*X*_, *α*_*G*_, *max*_*X*_, and *max*_*G*_, the tensor regression model will be fit to obtain 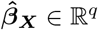 and 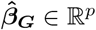. Due to the *L*^1^-norm truncating small values to zero from the elastic net constraint, only a subset of values within 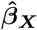 and 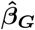 will be nonzero. Thus, final network features within data type ***𝒳*** will be the set 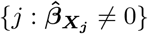, and final network features within data type ***𝒢*** will be the set 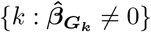. Algorithm 1 summarizes the steps to MOSCATO.

### 3.2 Model Tuning

MOSCATO assumes fixed values for *α*_*X*_, *α*_*G*_, *max*_*X*_, and *max*_*G*_ which will be tuned using an extension of the Stability Approach to Regularization Selection (StARS) method [22]. Tuning on accuracy, such as by cross validation or Bayesian information criterion, tends to result in overly dense solutions in high dimensional problems with results that are not reproducible [22]. The most extreme scenarios for stability are perfectly stable results from selecting no features (i.e., completely sparse) or selecting all features (i.e., no sparseness). Building on that logic, StARS initializes the parameters to the most sparse solution and gradually relaxes the sparsity until some instability threshold *ϕ* is met. Instability is estimated based on subsamples from the data by performing the feature selection under each subsample and summarizing the consistancy in results across different subsamples. Although *ϕ* may initially be thought of as an arbitrary cutoff between 0 and 1, it may be easily interpretted as the amount of allowable instability. In essence, a smaller *ϕ* would imply a more sparse but stable result. The motivation behind allowing some instability as opposed to fixing *ϕ* to 0 is to allow some noise to be selected in order to ensure that no true signal is missed in the final feature selection. In statistical terms, this means that StARS prioritizes reducing type II errors.

#### Algorithm 1 Schematic for MOSCATO

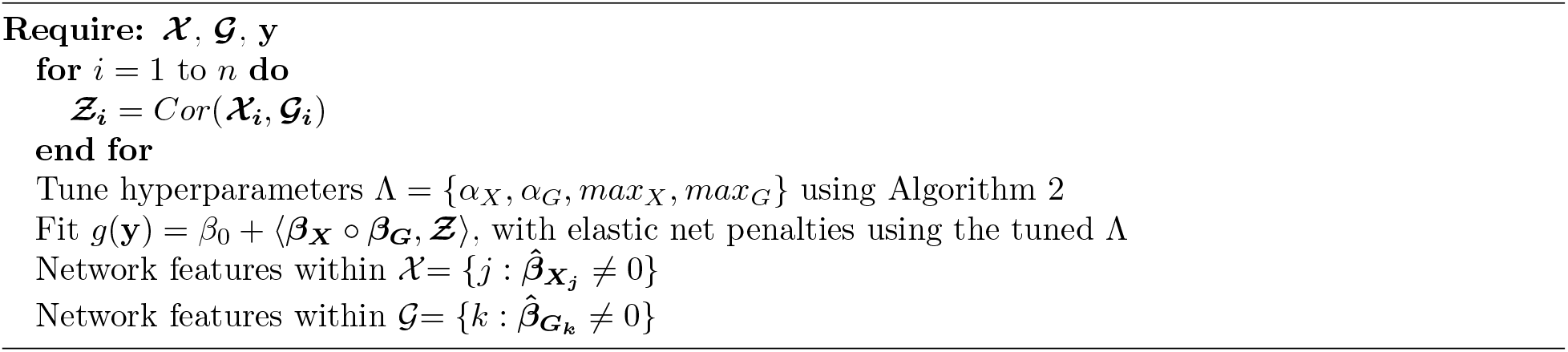

Although the StARS method was developed for tuning a single sparsity parameter, the four hyperparameters *α*_*X*_, *α*_*G*_, *max*_*X*_, and *max*_*G*_ will be tuned using similar logic. Focusing on tuning one dimension at a time, we initialize to a sparse solution with some small *max*_*X*_. Fixing *max*_*X*_, we estimate the instability for a range of *α*_*X*_ values between 0 and 1. Select the *α*_*X*_ value resulting in the lowest instability, and if that instability is less than *ϕ*, increase *max*_*X*_ and repeat the process. This continues until the *ϕ* instability is hit to select *max*_*X*_ and *α*_*X*_. Using the highest *max*_*X*_ and corresponding optimal *α*_*X*_ with instability less than *ϕ*, a similar process is then repeated for tuning *max*_*G*_ and *α*_*G*_. In this case with two dimensions, one for ***𝒳*** and another for ***𝒢***, we first tune *α*_*X*_ and *max*_*X*_ for some fixed *α*_*G*_ and *max*_*G*_, and then use 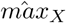 and 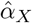 when tuning *α*_*G*_ and *max*_*G*_. The initial fixed *α*_*G*_ and *max*_*G*_ may be kept large, suppose *α*_*G*_ = 0.5 and *max*_*G*_ = *floor*(*p/*2) such that an overly sparse ***𝒢*** does not impact the stability on ***𝒳*** for the first dimension of tuning. This is summarized in Algorithm 2.

## 4 Simulations

To benchmark MOSCATO’s performance, it was applied to various simulations. The details on how the data was simulated is described in Section 4.1 and the results from MOSCATO are summarized in Section 4.2.

### 4.1 Simulation Details

Splatter [34] is a popular technique to simulate single-cell RNA-seq (scRNA-seq) data, and it has been shown to mimic distributions from real scRNA-seq data. The general Splatter schematic initiates by simulating a gene mean and then adjusts the gene mean to account for variation in outliers, library size, and dispersion. It then simulates the “observed” scRNA-seq values through a Poisson distribution using the adjusted gene mean, and the values are then randomly truncated to zero to replicate dropouts. Although Splatter realistically portrays scRNA-seq distributions, a few extensions were required in order to simulate multi-omic single-cell data with supervised gene networks.

To accomplish this, we leverage latent structures for multi-omic supervised networks detailed in Zhang et al. [35]. Figure 3 demonstrates the expected causal relationships within the data containing a supervised multi-omic network. In summary, we expect there to be a subgroup of features within ***𝒢*** and ***𝒳*** that relate to the outcome but not with each other; a subgroup of features within ***𝒢*** and ***𝒳*** that relate to each other but not with the outcome; a subgroup of features within ***𝒢*** and ***𝒳*** that are independent of each other and the outcome; and finally the target subgroups of the analysis, subgroup of network features within ***𝒢*** that relate to the network features within ***𝒳*** that ultimately relates to the outcome. These expected relationships among these latent components may be represented through a covariance matrix where the off diagonals will be non-zero where relationships exist and zero where independence is expected. More details are provided in the Appendix.

**Figure 3:**
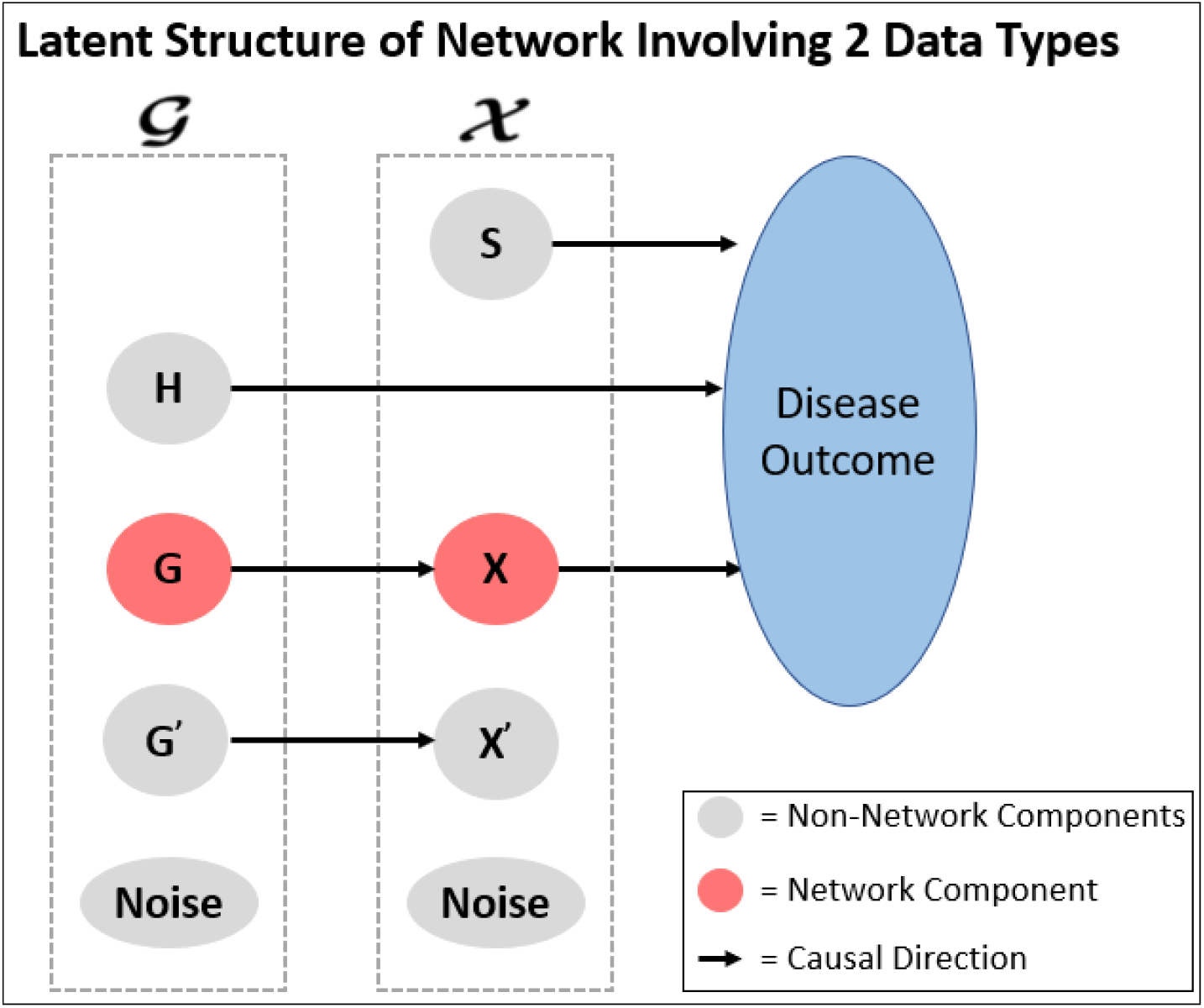
Latent structure for supervised multi-omic networks. From [30] and adapted from [35]. Assume two data types, ***𝒢*** and ***𝒳***. **S** describes the subgroup of features within ***𝒳*** that relate to the disease outcome but not with any features within ***𝒢***; **H** describes the subgroup of features within ***𝒢*** that relate to the disease outcome but not with any features within ***𝒳*** ; **G** and **X** describe the network where the group of features within ***𝒢*** relate to the group of features within ***𝒳*** that ultimately relate to the outcome; **G**′ and **X**′ denote the group of features that relate to each other but not with the outcome; and finally each data type will have independent noise not related to each other or the outcome.

#### Algorithm 2 StARS Method for Tuning Tensor Hyperparameter

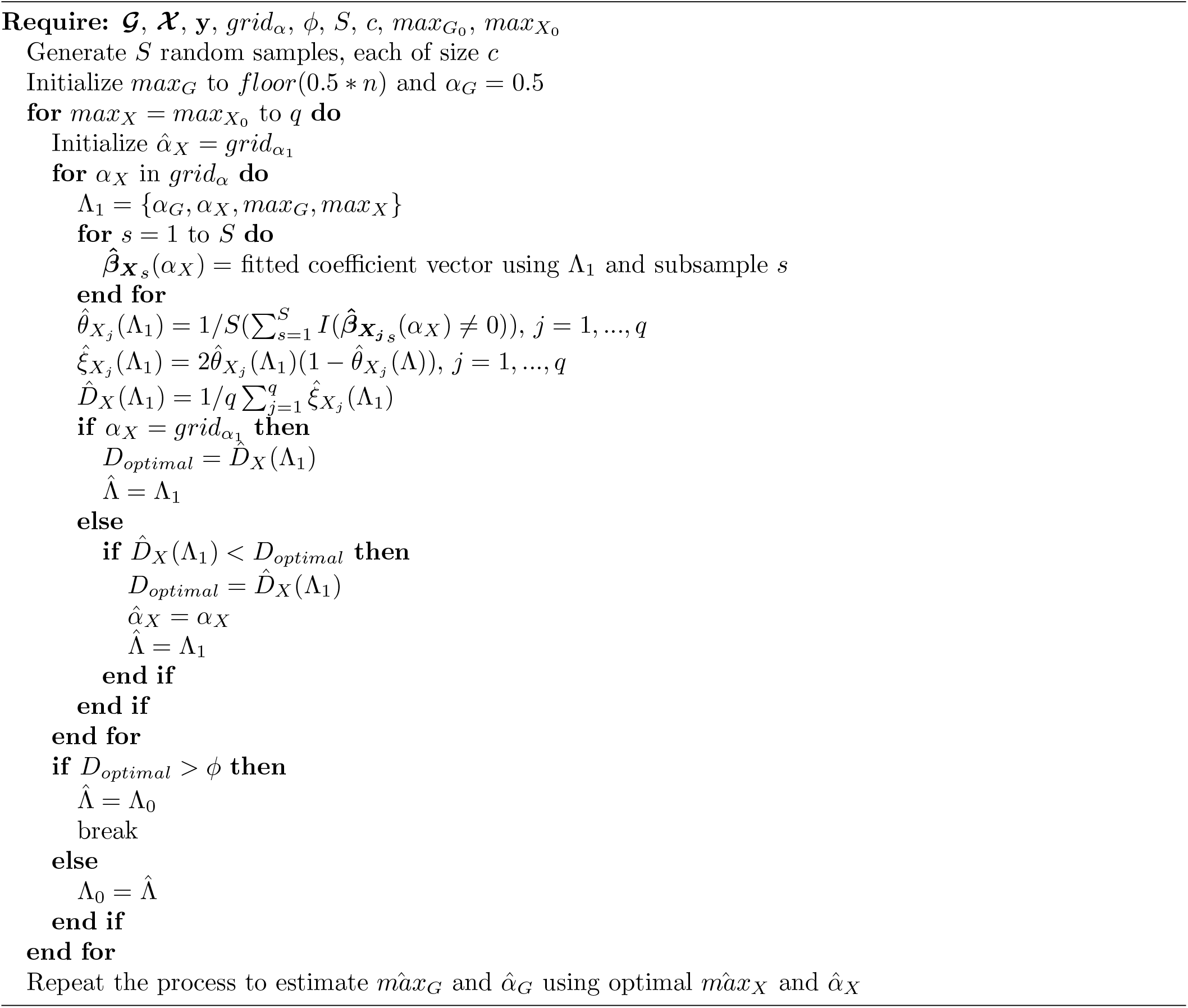

To extend Splatter for supervised multi-omic networks, we will first simulate the latent values (*S, H, X, G, G*′, *X*′, and *y*) for each subject using a multivariate normal distribution with zero mean and the latent variables’ covariance matrix. Since the latent values were simulated from a multivariate normal, they will each be normally distributed with mean 0. The Splatter simulation assumes the initial gene mean comes from a gamma distribution, so we take the square of the latent value divided by its standard deviation. By doing so, the transformed latent values then become gamma distributed with shape equal to 1/2 and scale equal to 2 times its variance. These transformed latent values will be used as the initial gene means for each subgroup of features, and the latent value for *y* will be used as the mean to simulate an observed outcome from a normal distribution. Initial gene means for the noise subgroup of features are randomly simulated independently. Additional theoretical details may be found in the Appendix.

Dispersion is adjusted on a subject level, library size is adjusted on a cellular level within a subject, outliers are adjusted on a feature level within a subject, and dropouts are accounted for on a cellular/feature level within a subject. The Splatter simulation is performed using the transformed gene means (combining all latent components within ***𝒢*** and ***𝒳***), and then the features are later separated by data type for each subject.

### 4.2 Results

MOSCATO was applied to a series of simulations using the techniques described in Section 4.1. Simulations were performed under 9 different settings accounting for the average number of cells per subject (250, 500, or 1000) and amount of technical noise (low, moderate, or high). Simulations were replicated 50 times under each simulation setting and each simulation had 100 subjects. ***𝒢*** contained 1440 total features where only 10 belonged to the network, and ***𝒳*** contained 1555 total features where only 15 belonged to the network. Table 1 describes the total number of features contained within each of the latent subgroups described in Figure 3.

**Table 1:** Number of Features in Simulations.

| Latent Set of Features | # Features |
| --- | --- |
| **H** | 15 |
| **G** | 10 |
| **G'** | 15 |
| Noise in $\mathcal{G}$ | 1400 |
| **S** | 20 |
| **X** | 15 |
| **X'** | 20 |
| Noise in $\mathcal{X}$ | 1500 |

The table summarizes the number of features within each latent subgroup used in the simulations. These latent subgroups are displayed in Figure 3. **H** describes the subgroup of features within *𝒢* that relate to the outcome but not any features within *𝒳*, **G** describes the subgroup of features within *𝒢* belonging to the network, **G**′ describes the subgroup of features within *𝒢* related to some features within *𝒳* but not the outcome, and **G**′ describes the subgroup of features within *𝒢* unrelated to features within *𝒳* and the outcome. **S, X, X**′, and Noise in *𝒢* describe subgroups of features within *𝒳* with similar relationships as the subgroups within *𝒢*.

For tuning the hyperparameters, we set *grid*_*α*_ = {0.2, 0.5, 0.7}, *ϕ* = 0.02, *R* = 50, and *c* = 50. 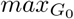 and 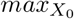 were initially set to 5, although this number may be increased in order to reduce runtimes as long as the instability remains below *ϕ* for its initialization. Ideally, MOSCATO would tune *max*_*G*_ to 10 and *max*_*X*_ to 15 in order to select the proper network size according to Table 1, but this will be unlikely due to the mechanics behind the StARS tuning method described in Section 3.2 which prioritizes reducing type II error over type I error. Figure 4 displays the tuned *max*_*G*_ and *max*_*X*_ across the simulations. As expected, all simulations tuned *max*_*G*_ and *max*_*X*_ to values greater than the true number of network features, regardless of the simulation setting. For *max*_*X*_, smaller values were tuned as technical noise decreased and number of cells increased, but this trend did not persist when tuning *max*_*G*_.

**Figure 4:**
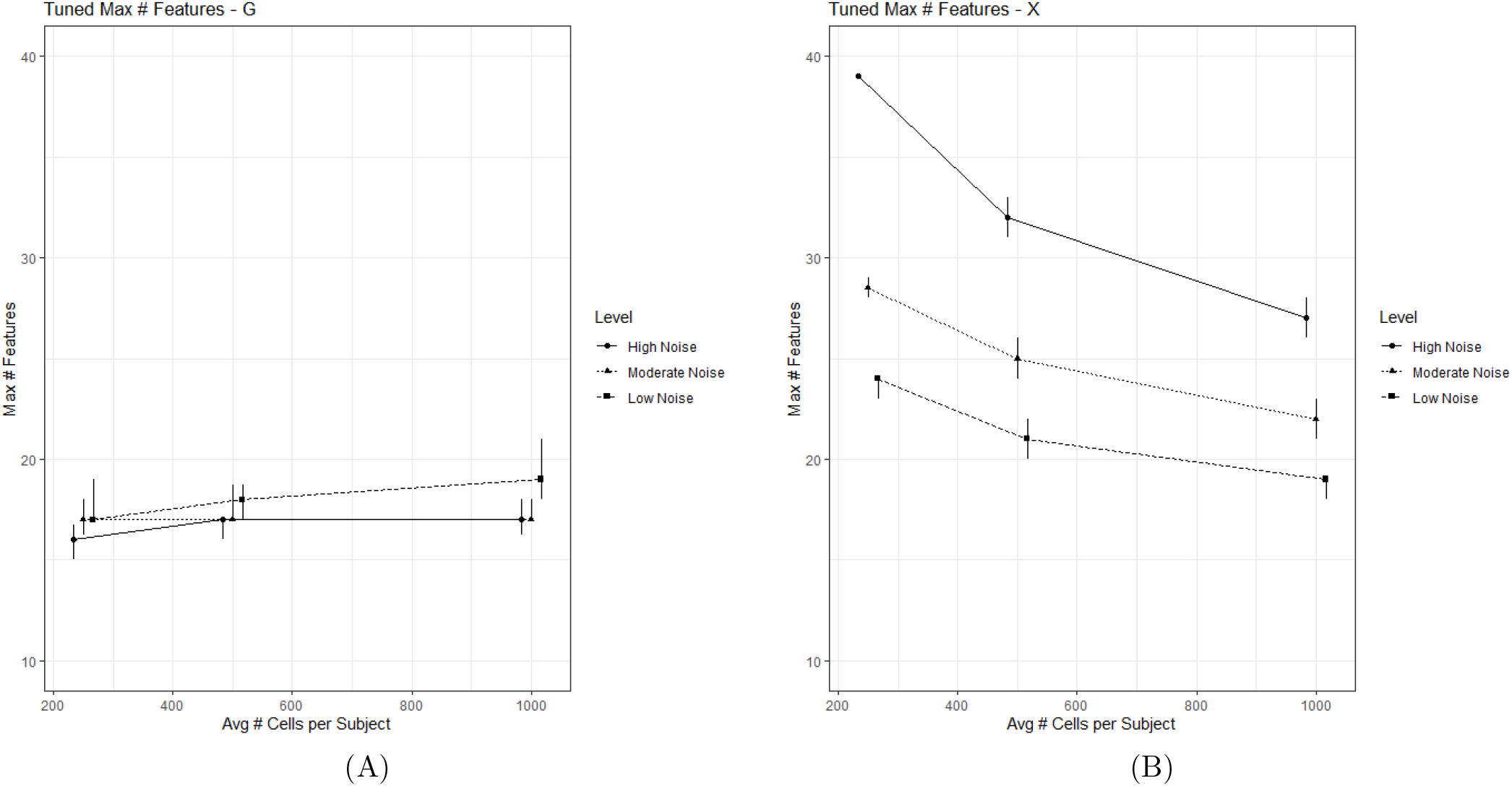
This figure displays the tuned *max*_*G*_ and *max*_*X*_ hyperparameters using the StARS method across the 9 different simulation settings accounting for different numbers of average cells per subject and different levels of technical noise. The points represent the median tuned values and the bars represent the first and third quartiles across 50 iterations for each simulation setting. (A) displays the tuned max network size within ***𝒢***, and (B) displays the tuned max network size within ***𝒳***. Ideally, MOSCATO would tune *max*_*G*_ to 10 and *max*_*X*_ to 15 in order to select the proper network size according to Table 1.

In addition to applying MOSCATO, we also applied competing methods using the area under the receiver operating curve (AUC). Seurat provides a popular single-cell sequencing workflow, and following similar methods used by the authors of Seurat [10], this AUC approach was done using the presto version 1.0.0 R package [19]. Selections using AUC were performed using two different criteria. One criteria was based on whether the Bonferroni adjusted p-value was less than the nominal significance level (set to 0.05) under the null hypothesis that the AUC equals 0.5. Additionally, selection criteria using cutoff values where features with an AUC either less than 0.3 or greater than 0.7 were selected. Since AUC requires categorical outcomes, we use the median of the outcome to binarize it (i.e., if the outcome is less than the median then recode the outcome as ‘0’, otherwise if the outcome is greater than the median then recode as ‘1’).

Figure 5 displays the sensitivity and specificity across the 9 simulation settings for data types ***𝒢*** and ***𝒳*** for the 3 different methods (MOSCATO, AUC using p-values, and AUC using cutoffs). Sensitivity measures the probability that network features are properly included in the selections, and specificity measures the probability that non-network features are properly excluded from the selections. As shown in Figure 5, the sensitivity and specificity under MOSCATO generally improve as the number of cells increases per subject and as technical noise decreases. Conversely, the specificity declines for AUC selections using p-values as the number of cells increases and the technical noise improves. AUC based on p-values not only produced counterintuitive results where the performance actually degraded as the technical noise reduced, it also selected too many features such that the results were not remotely sparse. This explains that while the sensitivity remained high for all simulations, this is simply due to the fact that nearly all features were selected using that criteria. AUC selections based on cutoffs resulted in opposite issues where it did not select nearly any features and produced poor sensitivity with nearly perfect specificity.

**Figure 5:**
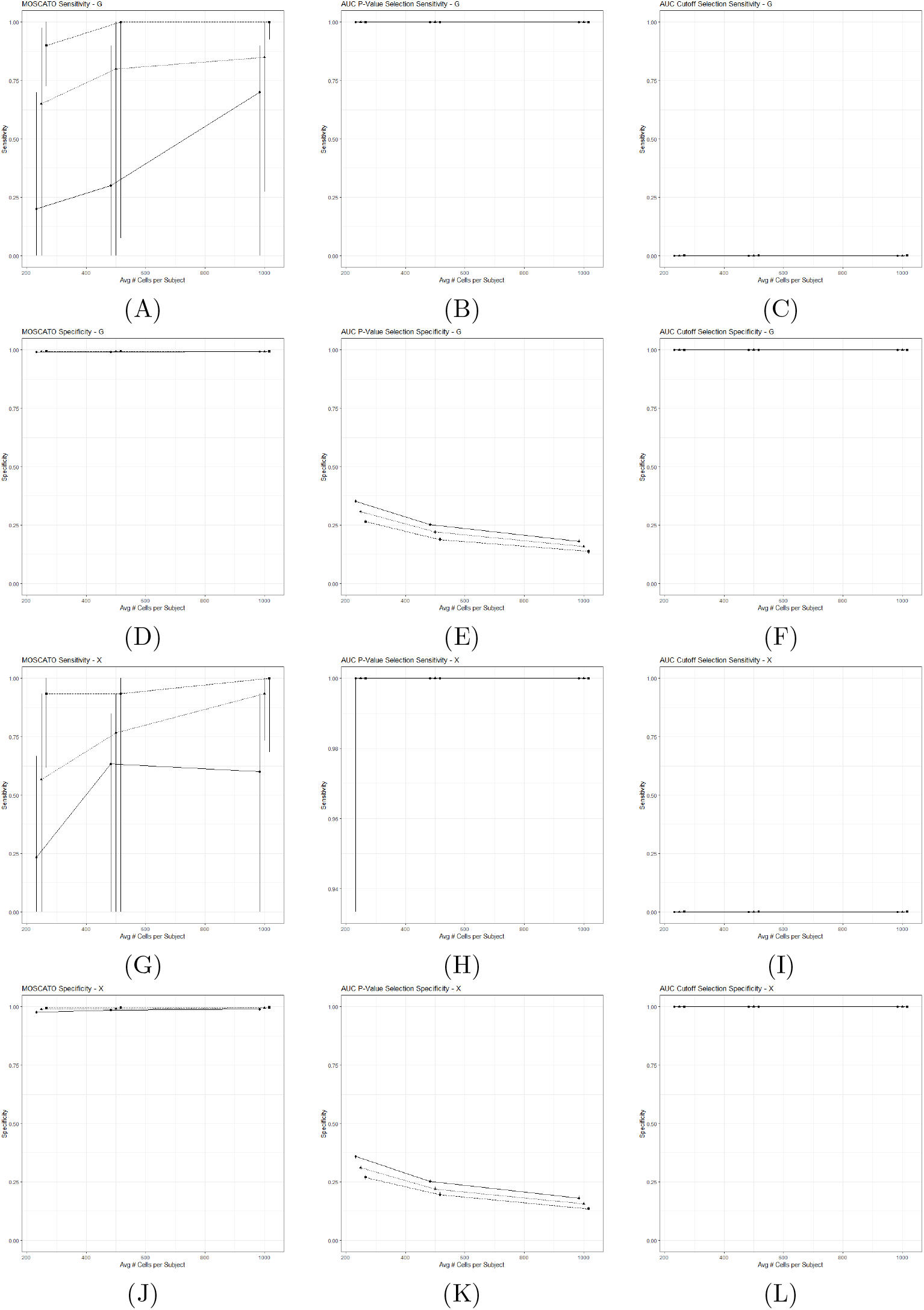
This figure displays the sensitivity and specificity across the 9 different simulation settings accounting for different numbers of average cells per subject and different levels of technical noise. The points represent the median sensitivity/specificity and the bars represent the first and third quartiles across 50 iterations for each simulation setting. (A)-(C) display the sensitivity for ***𝒢*** using MOSCATO, AUC selections using Bonferroni adjusted p-values, and AUC selections using cutoffs (< 0.3 or > 0.7). (D)-(F) display the specificity for ***𝒢*** under the three methods in the same order. Similarly, (G)-(I) displays the sensitivity for ***𝒳*** and (J)-(L) displays the specificity for ***𝒳***. Under perfect selections, the sensitivity and specificity should equal 1.

MOSCATO reproduced the superstructure of the network reasonably well with generally high sensitivity and also limited false positives present. This is especially true when comparing against approaches using the AUC. However, when it is expected that high levels of technical noise is present with limited cells per subject, caution should be used when considering MOSCATO.

## 5 Real Data Example

Leukemia encompasses all cancers that occur in blood cells. The 5-year survival rate is about 65% according to data from 2011 to 2017, and about 459,000 people were living with leukemia in 2018 in the United States [12]. Leukemia may be classified based on progression speed where chronic denotes slow progression and acute denotes aggressive progression. In addition to cancer progression, Leukemia may be subtyped by the type of cells where the cancer forms. For example, lymphocytic leukemia describes cancer developing from white blood cells and myelogenous leukemia describes cancer developing in blood forming cells within the bone marrow. Although rare, one may also have both lymphocytic and myelogenous leukemia which is denoted as mixed phenotype leukemia.

To assess MOSCATO in practice, we applied it to real single-cell data with multiple data types. Limited data is currently available due to the infancy of multi-omic single-cell sequencing, so data across multiple studies that all used CITE-seq protocols [29] on bone marrow / peripheral blood cells were used. CITE-seq produces cellular level RNA information and cell surface protein abundance (i.e., antibody derived tags (ADT)) simultaneously, and our outcome of interest will be leukemia versus healthy patients. Our goal will be to apply MOSCATO to this data in order to obtain a subset of RNA and ADT features associated with leukemia. After combining the data across studies, we have 14 healthy patients and 7 patients with leukemia. Of the 7 leukemia subjects, 1 had chronic lymphocytic leukemia (CLL) while the other 6 had mixed-phenotype acute leukemia (MPAL). The studies used to obtain the data are summarized in Table 2. Only non-perturbed, baseline cells were considered.

**Table 2:** Studies Used with CITE-seq Protocols and Healthy or Leukemia Subjects.

| Study ID | Tissue Type | # Healthy Subjects | # Leukemia Subjects |
| --- | --- | --- | --- |
| ERP124005 [21] | Blood | 10 | 0 |
| GSE152469 [4] | Blood | 0 | 1 |
| GSE139369 [9] | Blood | 1 | 4 |
| GSE139369 [9] | Bone Marrow | 3 | 2 |

Seurat version 4.0.3 [10] was used to normalize the data, cluster cells, and integrate the cell types across subjects. Figure 6 displays the Uniform Manifold Approximation and Projection (UMAP) [2] plots from the integrated data.

**Figure 6:**
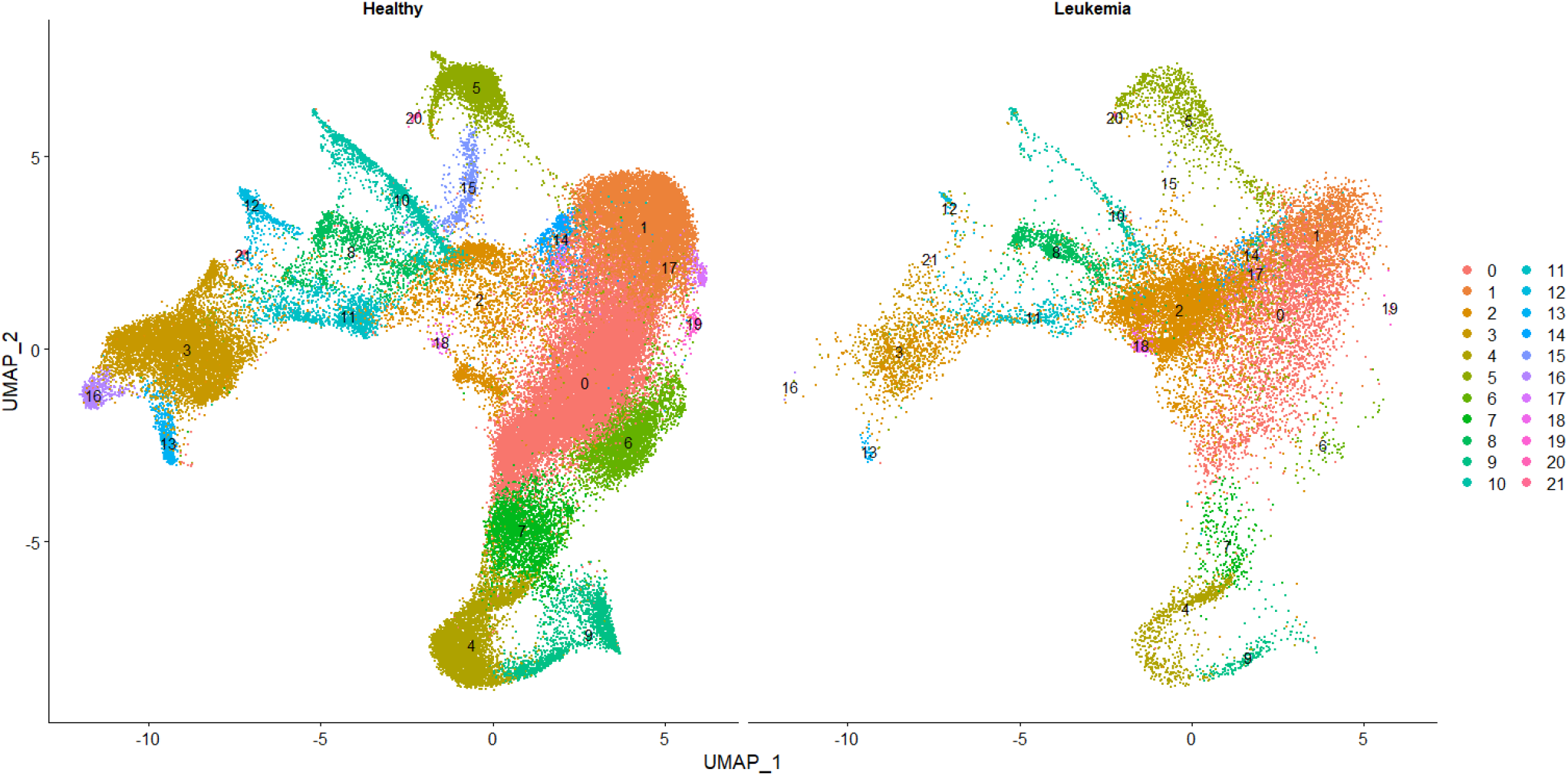
The UMAP results from the integrated cell types. The plots are split by healthy versus leukemia subjects.

After integration, 8 of the cell clusters (clusters 0, 1, 2, 3, 4, 5, 6 and 14 from Figure 6), 17991 RNA features, and 5 ADTs (CD3, CD4, CD14, CD19, and CD56) were measured across all subjects. We applied MOSCATO to each of these cell clusters separately, each with *grid*_*α*_ = {0.01, 0.05, 0.1}. Since Seurat clusters cells by maximizing correlation between features, the multicollinearity across features would consequently be high and require more weight on the *L*^2^-norm (i.e., lower *α* within the elastic net constraint).

Due to the modest sample size, we tuned the hyperparameters using a subsampling size of 20 (out of 21 total subjects) to estimate stability based on a “leave one out” scheme. Although StARS suggests setting the instability threshold *ϕ* to 0.05 for most applications [22], in this application with a small sample size (i.e., only 21 subsamples) and large number of RNA features (i.e., 17991 variables), the estimated instability under most sparse solutions will be much smaller than 0.05. For example, suppose all 21 subsamples select completely disjoint feature sets, but due to the high number of variables in consideration, many variables are consistently excluded from any selections in across all of the subsampled results. Since StARS considers both consistency in selections and consistency in exclusions, the estimated instability will be quite small due to the consistency in exclusions despite that the small number of features selected may be completely disjoint across all subsamples. Therefore, *ϕ* was set to 0.001.

To compare selections with another method, the MOSCATO results were compared to selections based on the AUC. The AUC approach was done using the presto version 1.0.0 R package [19]. Similarly as was done for MOSCATO, the AUC feature selections were performed on each cell cluster separately. Feature selections were made under two different selection criteria for AUC: if the Bonferroni adjusted p-value was less than 0.05 under the null hypothesis that the AUC equals 0.5 or whether the AUC was less tha 0.3 or greater than 0.7. Since the p-value would likely be small in situations with many cells (i.e., large sample sizes producing sensitive p-values for miniscule AUC deviations from 0.5), both a p-value approach and an approach based on the AUC values were considered.

### 5.1 Results

The complete results from MOSCATO, AUC selections based on p-values, and AUC selections based on the AUC cutoffs for each of the 8 cell clusters are provided in the supplementary files. In summary, the number of features selected by MOSCATO and AUC cutoffs were similarly sized, but AUC selections based on p-values resulted in nonsparse feature sets. DAVID [11, 28] was used to analyze and organize the gene ontology information from the RNA gene selections. DAVID clusters genes based on common annotations and functional information, and DAVID only allows clustering on gene sets with less than 3000 genes. Since the AUC selections based on p-values resulted in RNA selections well over this 3000 restriction for most cell clusters, we only focused on gene clusters from the MOSCATO and AUC cutoff selections.

MOSCATO selected 96 RNA features within cell cluster 0, and DAVID identified 2 gene clusters. The strongest gene cluster (based on highest enrichment score) used 7 of these 96 genes. This was the most notable gene cluster which included the genes CD3D, CD3E, and CD3G which are part of the KEGG pathway for Human T-cell Leukaemia Virus type 1 (HTLV-I) infection (KEGG pathway hsa05166), and HTLV-I infections are a known risk factor for developing adult T-cell leukaemia/lymphoma (ATL) [13]. Additionally, the genes CD2, CD3D, CD3E, CD3G, and CD4 within this gene cluster belong to the KEGG pathway for Hematopoietic cell lineage (hsa04640) which assists in producing blood cells. Given that the disease of interest in this application is based on leukemia (i.e., cancer in tissues which produce blood), it is reassuring that this gene cluster contains genes associated with blood production. Also, genes CD2, CD3D, CD3E, CD3G, KLRB1, CD247, and CD4 within the gene cluster are associated with the gene ontology for the cell surface receptor signaling pathway (GO:0007166) which makes sense given that MOSCATO summarized the single-cell data between RNA and cell surface proteins. Of the 5 cell surface proteins considered, MOSCATO selected the ADT’s CD3 and CD4 for this cell cluster. Figure 7 displays the RNA features within the strongest functional DAVID cluster, along with the ADT selections. No features selected by MOSCATO under cell cluster 0 were selected by AUC cutoffs, and although 83 of the 96 MOSCATO RNA selections were also selected by AUC based on p-values, the AUC selections based on p-values selected nearly half of all RNA features considered. AUC selections based on cutoffs selected 11 RNA features and 0 ADTs for cell cluster 0, and DAVID was not able to discern any gene clusters based on the genes selected.

**Figure 7:**
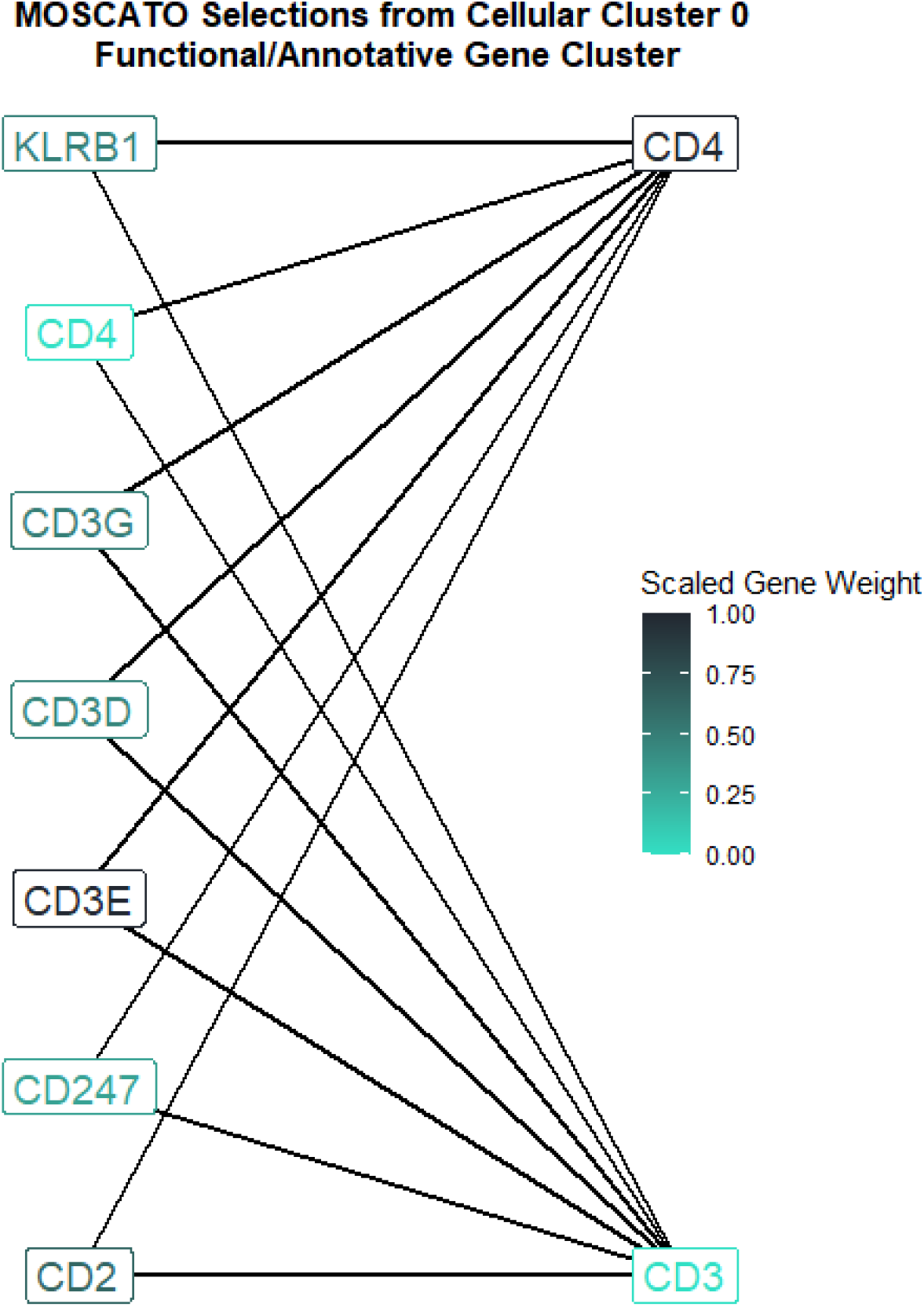
MOSCATO selected 96 RNA features and 2 ADT features from cell cluster 0 shown in Figure 6. This figure displays the strongest functional RNA gene cluster (left labels) within the 96 MOSCATO selections as determined by DAVID [11, 28], along with all ADT selections (right labels). DAVID determines gene clusters based on similar gene ontology and annotations. These RNA genes correspond to KEGG pathways for blood cell production (CD2, CD3D, CD3E, CD3G, and CD4) and HTLV-I infection (CD3D, CD3E, and CD3G), as well as genes with ontology information for cell surface receptor signaling (CD2, CD3D, CD3E, CD3G, KLRB1, CD247, and CD4). The label colors display the absolute scaled weight of the coefficient vectors from the tensor regression used in MOSCATO so that higher values correspond to a higher weight put on that gene/protein in the tensor regression.

In conclusion, the selections made by MOSCATO under cell cluster 0 resulted in a concise gene cluster discovered by previously known annotations and functionalities. These functionalities not only related to the disease of interest (i.e., leukemia), it also related to cell surface functionalities. This highlights that MOSCATO not only considers supervised information (e.g., disease versus no disease), it also considers the relationships across data types (e.g., RNA and cell surface proteins). Performing feature selection using solely AUC not only neglects the cross data type relationships, it also was not able to return a concise set of genes that related to leukemia. Furthermore, it is arbitrary to select pre-specified AUC cutoffs for selections, and the p-value selections did not produce sparse solutions. Also, AUC selections were based directly on normalized expression/sequenced values, but MOSCATO performs selections based on the similarities between data types. This possibly helps reduce batch effects found in individual subjects by standardizing each value between -1 and 1.

The real data application may be reproduced by following the steps provided at https://github.com/lorinmil/MOSCATOLeukemiaExample.

## 6 Discussion/Conclusions

MOSCATO was performed on both simulated and real data. The simulations produced fairly accurate results with a sensitivity and specificity close to 1 for many of the simulations, although MOSCATO did not perform as well in situations with high technical noise or low cell counts per subject. Since MOSCATO calculates its predictor tensor to be the estimated correlation of the datasets per individual, it is unsurprising that cell count would contribute to more accurate correlation estimates. Although not used in this manuscript, covariate adjustments could easily be made in MOSCATO by simply adding them to the tensor regression.

MOSCATO currently assumes all cells come from the same cell type, but it might be more interesting to accommodate situations in which multiple cell types are present. We are currently working on higher dimensional applications of MOSCATO in a future manuscript. A reasonable solution could incorporate another step which estimates a similarity matrix for each cell type and includes another dimension to the predictor tensor for cell type. Additionally, MOSCATO was only tested on experiments with 2 data types, but extensions should be considered in situations where more than two data types are present. This could be accomplished by extending the predictor tensor to accommodate a dimension per data type extracted, although higher dimensional summary measures would need to be considered.

MOSCATO was only tested using Pearson’s correlation as the summary measure to construct the predictor tensor 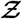, but other summaries should be considered. For example, mutual information or inverse covariance matrices might be interesting avenues to explore for future consideration. Graphical lasso [8] is a popular technique to obtain both the nodes and edges of a graph by applying lasso regressions to estimate the inverse of the covariance matrix, and this estimated inverse of the covariance matrix could be explored as the predictor tensor input for MOSCATO.

This study only used rank-1 tensors, but higher ranks could be considered that may unveil other patterns and networks available in the data. The proper rank could be obtained using typical model selection criteria such as cross validation, although interpreting the results may not be as straight forward.

Zhou et al. discuss hypothesis testing via asymptotic normality results [36], and these hypothesis testing schemes could be explored to assess network strength. For example, one could perform a global test whether the model coefficients equal zero for the network selections.

Although MOSCATO returns the superstructure for a graph, it does not provide information on directionality and does not currently consider directional consistency. For example, suppose two genes are positively correlated with another, but they contain opposing correlative directions with the outcome. This inconsistency in directionality makes interpreting the results more difficult, and may be mitigated by additionally tuning based on optimizing *balance*. This concept has been considered in bulk level analyses with a single omic type [31], and it could be considered for future work.

In summary, MOSCATO is a useful tool to indicate multi-omic features that relate to disease outcomes of interest to better accommodate multi-omic, single-cell data which is continuously growing in popularity.

## Supporting information

Real Data Selections

## 7 Availability

All code in this manuscript was done using R version 4.1.0 [27]. Code to perform MOSCATO and replicate the simulations may be found publicly on GitHub at https://github.com/lorinmil/MOSCATO. Steps and code to replicate the real data application may be found at https://github.com/lorinmil/MOSCATOLeukemiaExample.

