## Appendix for "MOSCATO: A Supervised Approach for Analyzing Multi-Omic Single-Cell Data"

September 2, 2021

### 1 Simulation Details

#### 1.1 Gene Means

Single-cell multi-omic data was simulated by extending the existing Splatter simulation scheme [2]. Splatter initially simulates the gene mean by randomly generating from a gamma distribution, but we instead will simulate the gene mean based on different latent relationships. These latent variables will be simulated separately for each subject based on a multivariate normal distribution with a common covariance matrix reflecting the dependencies. The latent variables include

- $y$: the latent variable for the outcome
- $X$: the latent variable for features within $\boldsymbol{\mathcal{X}}$ which relate to some features within $\boldsymbol{\mathcal{G}}$ as well with the outcome $y$
- $G$: the latent variable for features within $\boldsymbol{\mathcal{G}}$ which relate to some features within $\boldsymbol{\mathcal{X}}$ as well with the outcome $y$
- $S$: the latent variable for features within $\boldsymbol{\mathcal{X}}$ which relates to the outcome $y$ but not any features in $\boldsymbol{\mathcal{G}}$
- $H$: the latent variable for features within $\boldsymbol{\mathcal{G}}$ which relates to the outcome $y$ but not any features in $\boldsymbol{\mathcal{X}}$
- $X'$: the latent variable for features within $\boldsymbol{\mathcal{X}}$ that relate to some features within $\boldsymbol{\mathcal{G}}$ but not the outcome $y$
- $G'$: the latent variable for features within $\boldsymbol{\mathcal{G}}$ that relate to some features within $\boldsymbol{\mathcal{X}}$ but not the outcome $y$

Additionally we assume some group of features within $\boldsymbol{\mathcal{X}}$ and $\boldsymbol{\mathcal{G}}$ which do not relate to each other or the outcome, and we will call these features "noise". These relationships between latent variables were similarly used in other studies for bulk-level network analysis and similarly used here [3, 1]. These latent variable relationships are presented pictorally in Figure 3 in the manuscript.

Letting the latent variables be denoted as $\mathbf{L} = [y, X, G, S, H, X', G']^T$, we will set the expected value of $\mathbf{L}$ equal zero with covariance matrix

$$\boldsymbol{\Sigma} = \begin{bmatrix} \sigma_y^2 & \sigma_{y.x} & \sigma_{y.g} & \sigma_{y.s} & \sigma_{y.h} & 0 & 0 \\ \sigma_{y.x} & \sigma_x^2 & \sigma_{x.g} & 0 & 0 & 0 & 0 \\ \sigma_{y.g} & \sigma_{x.g} & \sigma_g^2 & 0 & 0 & 0 & 0 \\ \sigma_{y.s} & 0 & 0 & \sigma_s^2 & 0 & 0 & 0 \\ \sigma_{y.h} & 0 & 0 & 0 & \sigma_h^2 & 0 & 0 \\ 0 & 0 & 0 & 0 & 0 & \sigma_{x'}^2 & \sigma_{x'.g'} \\ 0 & 0 & 0 & 0 & 0 & \sigma_{x'.g'} & \sigma_{g'}^2 \end{bmatrix} \tag{1}$$

For each subject $i$, their latent variables $\mathbf{L}_i$ will be randomly generated from $N_7(\mathbf{0}, \mathbf{\Sigma})$. Since the Splatter simulations assume the gene means come from some gamma distribution, we transform the generated latent variables into a gamma distribution by first dividing its elements by their standard deviations (i.e., the square root of the diagonal elements within $\mathbf{\Sigma}$) and then squaring. This transformation from multivariate normal to marginal gamma distributions is detailed in the proof for Theorem 1.1,

**Theorem 1.1** *If $\boldsymbol{L} \sim N_7(\boldsymbol{0}, \mathbf{\Sigma})$ then $L_j^* = (L_j/\sqrt{\Sigma_{jj}})^2$ is distributed as $\Gamma(1/2, 2)$, where $L_j$ denotes the $j^{th}$ element in $\boldsymbol{L}$ and $\Sigma_{jj}$ denotes the $j^{th}$ diagonal element in $\mathbf{\Sigma}$.*

**Proof 1.1** *Since $\boldsymbol{L} \sim N_7(\boldsymbol{0}, \mathbf{\Sigma})$ is multivariate normal, then the elements $L_j$ are marginally distributed as $N(0, \Sigma_{jj})$. Then, it may be transformed into standard normal by $L_j/\sqrt{\Sigma_{jj}} \sim N(0,1)$. Finally, $(L_j/\sqrt{\Sigma_{jj}})^2$ transforms into a $\chi_1^2$ distribution because it is the square of a standard normal, and $\chi_1^2$ is equivalent to $\Gamma(1/2, 2)$.*

Letting this gamma transformed latent variable be denoted as $\mathbf{L}_i^*$, we now have gene means that may be used for Splatter. For example, for the features belonging to the subgroup of features within $\mathbf{X}$, we would use the second element in $\mathbf{L}_i^*$ as the gene mean and continue through the typical Splatter schematic to generate its "observed" values across the cells. Similar logic would hold for the other features belonging to the different gene groups by using the transformed simulated latent variables as the means.

The gene means for the noise features were generated by first generating its value from a normal distribution with mean 0 and variance 1, and then square the value to get the gene mean. This is repeated independently for each of the noise features within $\boldsymbol{\mathcal{X}}$ and $\boldsymbol{\mathcal{G}}$.

The "observed" univariate outcome for a given subject $i$ was generated from a normal distribution using the first element within $\mathbf{L}_i^*$ as the mean and $\tau\sigma_y^2$ as the variance where $\tau$ may be used to adjust the network signal strength.

#### 1.2 Multi-Omics using Splatter

Splatter was created to simulate single-cell RNA-seq data, but now we want to simulate two different data types. To accomplish this, any cellular-level simulated values from Splatter which were used to modify the gene mean were held equal between $\boldsymbol{\mathcal{X}}$ and $\boldsymbol{\mathcal{G}}$. Since multi-omic single-cell data is done on a simultaneous level, this ensures that the cells are related across data types in some way. Any feature-level modifications done to the gene mean within Splatter were kept on an individual feature level. In summary, the gene means are extracted using the latent information, the remainder of the Splatter schematic still holds, and then the data types are separated into two different datasets. Additionally, although the number of cells was not addressed in Splatter, the number of cells for a subject $i$ was randomly generated from a Poisson distribution.

#### 1.3 Simulation Parameter Values

Simulations were performed under 9 different simulation settings: 3 different average number of cells per subject (250, 500, and 1000) and 3 different technical noise levels (low, moderate, high). $\mathbf{\Sigma}$ was held constant across all simulation settings to equal

$$\begin{bmatrix} 1.4180796 & 0.6281485 & 1 & 0.1441176 & 0.35 & 0 & 0 \\ 0.6281485 & 0.4117647 & 0.591608 & 0 & 0 & 0 & 0 \\ 1 & 0.591608 & 1 & 0 & 0 & 0 & 0 \\ 0.1441176 & 0 & 0 & 0.4117647 & 0 & 0 & 0 \\ 0.35 & 0 & 0 & 0 & 1 & 0 & 0 \\ 0 & 0 & 0 & 0 & 0 & 0.4117647 & 0.591608 \\ 0 & 0 & 0 & 0 & 0 & 0.591608 & 1 \end{bmatrix} \quad (2)$$

and $\tau = 0.384083$. For the technical noise levels, the values for the Splatter simulation are summarized in Table 1.

Table 1: **Technical Noise Levels in Simulations**

| Description | Parameter | Low Noise | Moderate Noise | High Noise |
|---|---|---|---|---|
| Outlier Probability | $\pi_0$ | 0.002 | 0.01 | 0.05 |
| Outlier Location | $\mu_0$ | 5 | 5 | 5 |
| Outlier Scale | $\sigma_0$ | 0.4 | 0.4 | 0.4 |
| Library Size Location | $\mu_L$ | 12 | 12 | 12 |
| Library Size Scale | $\sigma_L$ | 0.2 | 0.2 | 0.2 |
| Common Dispersion | $\phi$ | 0.1 | 0.1 | 0.1 |
| BCV df | $df_{BCV}$ | 7 | 7 | 7 |
| Dropout Midpoint | $x_0$ | 1 | 1.5 | 2 |
| Dropout Shape | $k_{shape}$ | 0.5 | 0.4 | 0.3 |

### 2 Real Data Application

#### 2.1 Additional Information on Feature Selections

Table 2 summarizes the selections made by MOSCATO and AUC. Note that two different selection criteria was used for AUC. The Bonferroni adjusted p-value being less than the nominal significance level (set to 0.05) or based on whether the AUC was less than 0.3 or greater than 0.70. For all cell clusters, selecting features based on the AUC p-value resulted in much larger selections than based solely on the AUC estimate or MOSCATO. In fact, the AUC's p-value selected over half of the featues for Cluster 1 and almost half of the features for Cluster 0.

Table 2: **Size of RNA Feature Selections**

| **Cell Cluster** | **MOSCATO** | **AUC ($p - value < 0.05$)** | **AUC ($AUC < 0.3$ or $> 0.7$)** |
|---|---|---|---|
| 0 | RNA: 96, ADT: 2 | RNA: 8473, ADT: 5 | RNA: 11, ADT: 0 |
| 1 | RNA: 10, ADT: 2 | RNA: 9393, ADT: 5 | RNA: 37, ADT: 2 |
| 2 | RNA: 40, ADT: 5 | RNA: 6321, ADT: 5 | RNA: 49, ADT: 4 |
| 3 | RNA: 20, ADT: 1 | RNA: 2917, ADT: 5 | RNA: 58, ADT: 5 |
| 4 | RNA: 17, ADT: 2 | RNA: 3963, ADT: 5 | RNA: 32, ADT: 4 |
| 5 | RNA: 20, ADT: 2 | RNA: 1632, ADT: 5 | RNA: 8, ADT: 3 |
| 6 | RNA: 8, ADT: 3 | RNA: 1355, ADT: 4 | RNA: 29, ADT: 3 |
| 14 | RNA: 40, ADT: 5 | RNA: 3579, ADT: 5 | RNA: 129, ADT: 5 |

The table displays the number of features selected by the 3 different methods. MOSCATO selects features by utilizing tensor regression based on each subject's single-cell multi-omic data (involving RNA and ADT features), AUC ($p - value < 0.05$) selects features by Bonferroni adjusted p-values for testing whether the AUC equals 0.5 from cells of leukemia subjects versus cells of healthy subjects, and AUC ($AUC < 0.3$ or $> 0.7$) selects features whose AUC is either less than 0.3 or greater than 0.7. The data contains 17991 total RNA features and 5 ADT features.

The RNA selections made from each of the cell clusters were input into DAVID. DAVID only found functional gene clusters for cell cluster 0.
